# A genomic resource for exploring bacterial-viral dynamics in seagrass ecosystems

**DOI:** 10.1101/2024.12.06.627215

**Authors:** Cassandra L. Ettinger, Jason E. Stajich

## Abstract

**Background:** Seagrasses are globally distributed marine flowering plants that play foundational roles in coastal environments as ecosystem engineers. While research efforts have explored various aspects of seagrass-associated microbial communities, including describing the diversity of bacteria, fungi and microbial eukaryotes, little is known about viral diversity in these communities.

**Results:** To begin to address this, we leveraged metagenomic sequencing data to generate a catalog of bacterial metagenome-assembled genomes (MAGs) and phage genomes from the leaves of the seagrass, *Zostera marina*. We expanded the robustness of this viral catalog by incorporating publicly available metagenomic data from seagrass ecosystems. The final MAG set represents 85 high-quality draft and 62 medium-quality draft bacterial genomes. While the viral catalog represents 354 medium-quality, high-quality, and complete viral genomes. Predicted auxiliary metabolic genes in the final viral catalog had putative annotations largely related to carbon utilization, suggesting a possible role for phage in carbon cycling in seagrass ecosystems.

**Conclusions:** These genomic resources provide initial insight into bacterial-viral interactions in seagrass meadows and are a foundation on which to further explore these critical interkingdom interactions. These catalogs highlight a possible role for viruses in carbon cycling in seagrass beds which may have important implications for blue carbon management and climate change mitigation.

## Background

Seagrasses are submerged marine flowering plants that have critical roles as foundation species in coastal ecosystems worldwide. They provide essential ecosystem services such as stabilizing the seafloor, filtering pollutants, supporting fisheries, and driving biogeochemical cycles [1–3]. One of their most significant contributions is their ability to sequester carbon in both their tissues and surrounding sediments (i.e., blue carbon) [2,4]. Despite their ecological importance, seagrass ecosystems are under increasing threat from pollution, climate change, and coastal development. Preserving these ecosystems is critical not only for their carbon sequestration potential but also for maintaining the biodiversity and ecological services they provide to coastal communities.

In recent years, there has been growing recognition of the importance of microorganisms in maintaining the health of plants [5–7]. Studies have begun to describe the composition and structure of the bacterial community associated with seagrasses, particularly *Zostera marina*, the dominant seagrass in the Northern hemisphere [8–14]. These bacterial communities are thought to have important roles in facilitating nitrogen and sulfur cycling to benefit seagrass growth and survival [8,9,11,12,15–17].

While research has largely focused on the bacterial component of these microbial communities, advances in sequencing and bioinformatics now allow us to explore the role of viruses in these systems. Phages, in particular, have been shown to influence global ocean and soil biogeochemical cycles by modulating host population dynamics through auxiliary metabolic genes (AMGs), which can directly alter bacterial metabolism to increase overall host fitness [18–21]. However, the diversity and ecological roles of phage in seagrass ecosystems remains largely unexplored.

In this study, we generated a catalog of viruses from seagrass-associated metagenomic samples, and then investigated host-viral dynamics of viral operational taxonomic units (vOTUs) and bacterial metagenome-assembled genomes (MAGs) generated from *Z. marina* leaf samples from Bodega Bay, CA. Specifically our objectives were to: (i) create a catalog of vOTUs from *Z. marina* and other seagrass species using publicly available metagenomic sequencing data, (ii) assemble a catalog of bacterial MAGs from *Z. marina* leaf tissue, and (iii) explore bacterial-phage dynamics with a focus on exploring AMGs involved in nitrogen and sulfur metabolisms.

## Methods

### Sequence generation

We extracted DNA from epiphytic washes from *Z. marina* leaves as part of previous work focused on characterizing the mycobiome using high-throughput sequencing of the ITS2 region [22]. We chose three DNA extracts from that work here for deep metagenomic sequencing with the goal of obtaining high quality metagenome-assembled genomes. We provided DNA to the UC Davis Genome Center DNA Technologies Core for sequencing and library preparation. DNA libraries were sequenced on an Illumina HiSeq4000 to generate 150 bp paired-end reads.

### Metagenomic processing

We trimmed sequence reads using bbDuk v. 37.68 [23] with the following parameters: qtrim=rl trimq=10 maq=10. We then mapped against and removed any reads from the metagenomes matching the available genome for *Z. marina* v. 3.1 [24] using bowtie2 v. 2.4.5 [25] and samtools v. 1.11 [26]. We co-assembled the remaining reads from all three metagenomic samples using MEGAHIT v. 1.2.9 [27].

We identified and assessed metagenome-assembled genomes (MAGs) using the anvi’o v. 7.2 workflow [28]. First, we used bowtie2 v. 2.4.5 [25] and samtools v. 1.11 [26] to obtain read coverage for each metagenomic sample against the assembly. Then we used “anvi-gen-contigs-database” to generate a database from the co-assembly and to predict open-reading frames using Prodigal v. 2.6.3 [29]. This command also identifies single-copy bacterial [30], archaeal [31], and protista [32] genes using HMMER v. 3.2.1 [33] and ribosomal RNA genes using barrnap [34]. We predicted taxonomic assignments for each gene call using Kaiju v. 1.8.2 [35] with the NCBI BLAST non redundant protein database nr including fungi and microbial eukaryotes v. v. 2020-05-25. For each individual metagenomic sample, we then used “anvi-profile” to construct an anvi’o profile for contigs >1 kbp with the “–cluster-contigs” option. Next we ran several automatic binning algorithms including MetaBAT2 v. 2.15, MaxBin v. 2.2.1, BinSanity v.0.5.4 and CONCOCT v. 1.1.0 [36–39] to generate preliminary sets of bacterial MAGs. We provided the resulting set from each algorithm to DAStool v. 1.1.2 [40] to generate a single optimal MAG set. MAGs from this set were then manually assessed for contamination and refined using “anvi-refine”. After manual refinement, MAGs were further de-contaminated using MAGpurify v. 2.1.2 [41].

We assessed MAG completeness and contamination using CheckM v. 1.2.1 [42] and CheckM2 v. 1.0.1 [43]. In this work we report all identified MAGs with >80% completion and <10% contamination. We refer to MAGs as high quality if they were >90% complete with <5% contamination, and medium quality if they were >50% complete with <10% contamination per established guidelines [44]. To obtain a putative taxonomy for each MAG, we used GTDB-Tk v. 2.2.6 [45], which uses a combination of average nucleotide identity and phylogenetic placement in the context of the Genome Taxonomy Database to taxonomically identify MAGs.

### Viral identification

In order to have a robust dataset to compare our recovered virus catalog in the context of other studies, we downloaded publicly available data from NCBI GenBank from nine studies [11,12,17,46–51] representing 65 metagenomic and 18 metatranscriptomic samples collected from seagrass environments (Table S1).

We filter and trimmed public metagenomic data using bbDuk v. 37.68 [23] with the following parameters: ktrim=r k=23 mink=11 hdist=1 tpe tbo qtrim=rl trimq=10 maq=10. Then for each study, samples were co-assembled using MEGAHIT v. 1.2.9. We used the co-assemblies for each study, as well as the co-assembly from this study, when identifying viral sequences using a workflow similar to Guo et al. [52]

Briefly, we identified viral sequences from metagenomic co-assemblies using VirSorter2 v. 2.2.3 [53], a tool that uses multiple random forest classifiers to predict whether a sequence contains a DNA or RNA virus, with the following parameters: --min-length 5000, --min-score 0.5, --include-groups dsDNAphage,RNA,ssDNA,lavidaviridae. We then ran CheckV v0.8.1 [54] on the VirSorter2 predicted viral sequences using the “end_to_end” workflow. To be conservative in our analyses, we removed viral sequences with a CheckV quality score of “not-determined” and “low-quality” prior to downstream analysis. We then ran VirSorter2 again on the viral sequences from the CheckV workflow with the --prep-for-dramv option. We used DRAM-v v. 1.2.2 [55] to “annotate” viral sequences and then “distill” annotations into predicted auxiliary metabolic genes (AMGs) for phage.

### Virus clustering and analysis

We clustered viral sequences ≥ 10 kbp in length into 95% similarity viral operational taxonomic units (vOTUs) using dRep v. 3.2.2 [56]. We used Prodigal v. 2.6.3 [29] to predict open reading frames in vOTUs using the -p meta option. We then provided the predicted proteins from the phage vOTUs to VContact2 v. 0.9.19, as well as predicted proteins from the INPHARED August 2023 viral reference database, to generate viral clusters (VCs) based on viral gene-sharing networks [57,58]. We further used geNomad v. 1.7.4 to assign taxonomy to phage vOTUs [59]. We used iPHoP v. 1.3.2 [60], which integrates across multiple methods in a machine learning framework to assign host taxonomy at the genus level, to predict host-virus linkages using a combination of the final bacterial MAG collection from this study and iPHoP’s reference host database.

We mapped reads from each metagenome to vOTUs using bowtie2 --sensitive with a minid=0.90 to quantify vOTU relative abundance [61]. We then used SAMtools and BEDTools genomecov to obtain coverage estimates for each vOTU across each individual metagenomic sample [26,62]. We used coverM in contig mode to parse bam files and calculate the trimmed pileup coverage (tpmean) of vOTUs which displayed ≥ 75% coverage over the length of the viral sequence. Thresholds for analysis of vOTUs were based on community guidelines for length (i.e. ≥ 10 kbp), similarity (i.e. ≥ 95% similarity), and detection (i.e. ≥ 75% of the viral genome length covered ≥ 1x by reads at ≥ 90% average nucleotide identity) [63,64]. The viral relative abundance (tpmean), CheckV quality, geNomad taxonomy, iPHoP host-prediction, MAG diversity, and DRAM-v annotation results were analyzed and visualized in R v. 4.3.0 [65] using the tidyverse v. 2.0.0 and phyloseq v. 1.44.0 [66,67].

To compare viral diversity between metagenomic samples (i.e. beta diversity), we calculated the Hellinger distance, the Euclidean distance of Hellinger transformed relative abundance (tpmean) data. We performed Hellinger transformations using the transform function in the microbiome v. 1.22.0 package [68], calculated the Hellinger distance using the ordinate function in phyloseq, and then visualized these distances using principal-coordinate analysis (PCoA).

## Results and Discussion

### Refined viral catalog from seagrass ecosystems

In total, we recovered 17,145 predicted viral genomic fragments which we clustered at 95% average nucleotide identity into 3,633 vOTUs. To ensure a high quality viral catalog, we filtered this dataset further using community thresholds for length and quality [63,64]. The refined viral collection represents 354 viral sequences comprising 351 double-stranded DNA phage, two lavidaviridae, and one RNA phage based on VirSorter2 random forest classification, with each sequence representing a unique vOTU (Table S2). Of these sequences, nine are integrated prophage (Figure 1A). The final catalog includes 44 complete, 28 high-quality and 282 medium-quality draft viral genomic fragments (Figure 1B).

**Figure 1.**
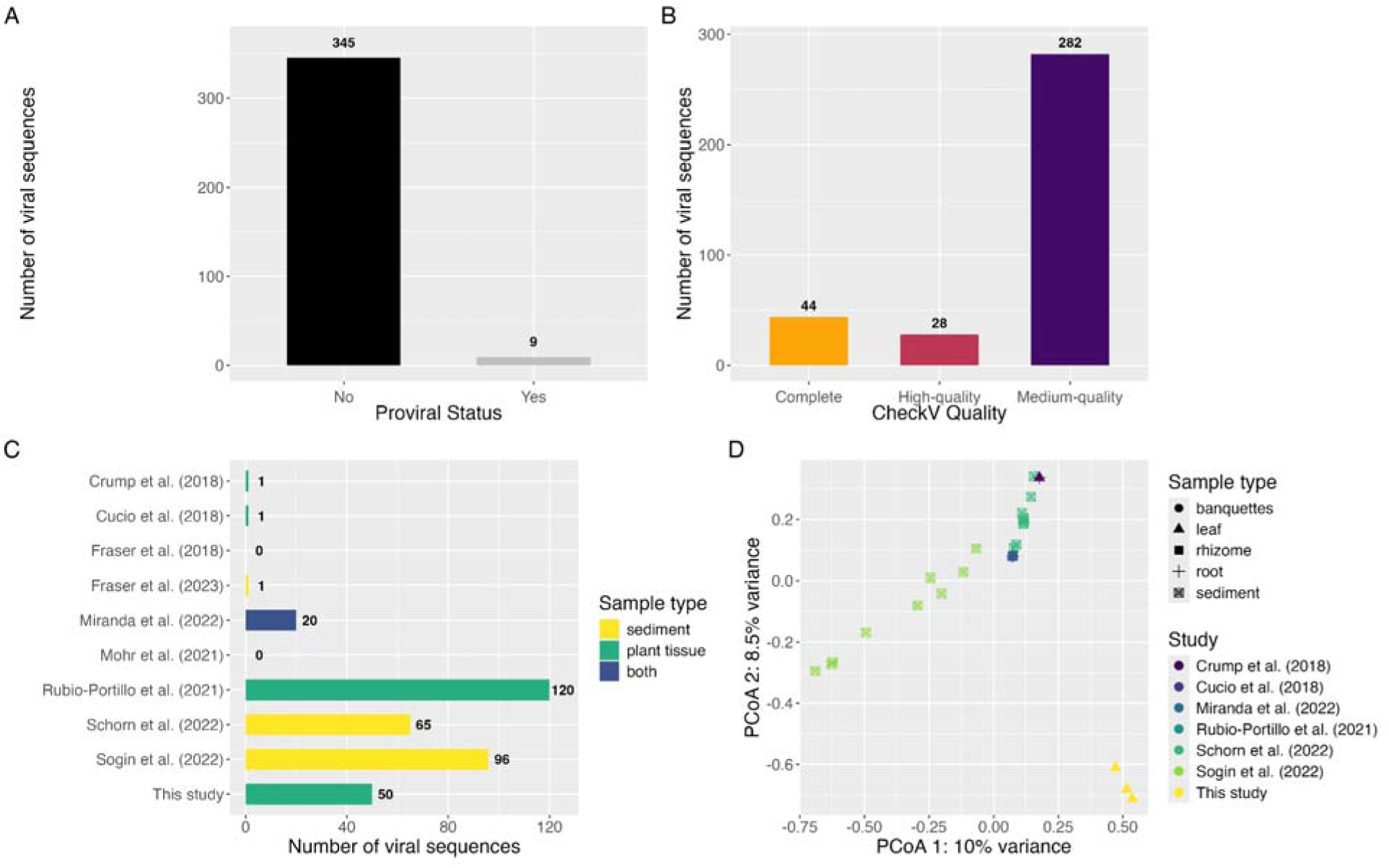
Putative viral sequences identified from seagrass ecosystems. (A) The proviral status of predicted viral genomes is shown as a bar graph, with the number above the bar representing the total number of sequences. (B) CheckV quality metrics for viral sequences are shown as a bar graph with bars colored by quality and the number above the bar representing the total number of sequences in each bar. (C) Bar chart depicting the number of viral sequences identified in the metagenomic co-assembly from each study, with the number to the right of the bar representing the total number of sequences in each bar. (D) Principal-coordinate analysis (PCoA) visualization of Hellinger distances of relative abundance of viral communities across metagenomic samples. Samples are colored by study, and have shapes based on sample type.

To explore the taxonomy of the viral sequences reported here, we used two complementary approaches: VContact2 [57], a cluster based method, and geNomad [59], an alignment based tool. We ran VContact2 [57] with INPHARED [58] reference genomes to cluster phage vOTUs into 136 VCs, which represent genus-level groupings based on gene-sharing networks. Of these, 35 VCs represented clusters that contained reference genomes, while the other 101 VCs were unique, potentially representing novel phage genera (Figure S1A). However, 148 vOTUs could not be assigned confidently to any VC.

Taxonomic classification of viruses was further refined using geNomad [59]. Most vOTUs (98.58% of vOTUs) were assigned to the *Caudoviricetes* class of tailed double-stranded DNA bacteriophage (Figure S1B). However, viral taxonomy is currently in revision [69], and the majority of sequences could not be confidently placed into finer taxonomic ranks: 331 vOTUs were unclassified at the order level, and 352 at the family level. While geNomad provided higher-level taxonomic predictions for 351 of the 354 vOTUs, more comprehensive classification awaits future updates to formal viral taxonomy definitions.

The recovery of phage genomes from metagenomes varied across experimental studies (Figure 1C). Of the 354 viral sequences in the final catalog, 50 were recovered from the new metagenomic data in this study, with the remainder derived from publicly available metagenomes. Studies that contributed more viral sequences to the final catalog generally employed deeper sequencing, with average depths ranging from 9.7 to 28.2 Gb per sample, compared to studies with few recovered viral sequences (1.2 to 9.5 Gb per sample).

When we examined viral community composition across metagenomes, clustering by scientific study was evident (Figure 1D). Notably, the viral communities in the metagenomes generated in this study formed a distinct cluster. However, technical differences (e.g., sequencing depth) across datasets prevented us from drawing broader conclusions regarding viral diversity based on seagrass species or tissue type.

Despite the use of deep metagenomic sequencing and the integration of public metagenomes, we recovered a relatively small, refined catalog of viral sequences. This highlights the limitations of relying solely on short-read metagenomic sequencing to capture viral diversity, at least in seagrass ecosystems. Viromics approaches (e.g., [70]) may provide more comprehensive insights into these viral communities, as even with improved bioinformatics pipelines, true viral diversity and abundance remains elusive.

### MAG collection reflects abundant bacterial groups on Z. marina leaves

We assembled 147 total bacterial MAGs including 85 high-quality draft MAGs ( >90% completion, <5% contamination), and 62 medium-quality MAGs (>80% completion, <10% contamination) (Table S3). These MAGs largely belong to the following taxonomic classes (Figure 2A): Alphaproteobacteria (43), Gammaproteobacteria (35), Bacteroidia (29), and Plactomycetia (16). Within these classes, the most frequently recovered orders were: Rhodobacterales (28), Pirellulales (14), Flavobacteriales (12), Chitinophagales (11) and Pseudomonadales (11). Notably, using GTDB-Tk 30 MAGs could not be assigned to any known genus, six were unclassified at the family level, and one lacked an assignment at the order level, highlighting potential evolutionary novelty compared to the current GTDB database. These findings suggest that seagrass ecosystems may harbor novel bacterial species that remain uncharacterized.

**Figure 2.**
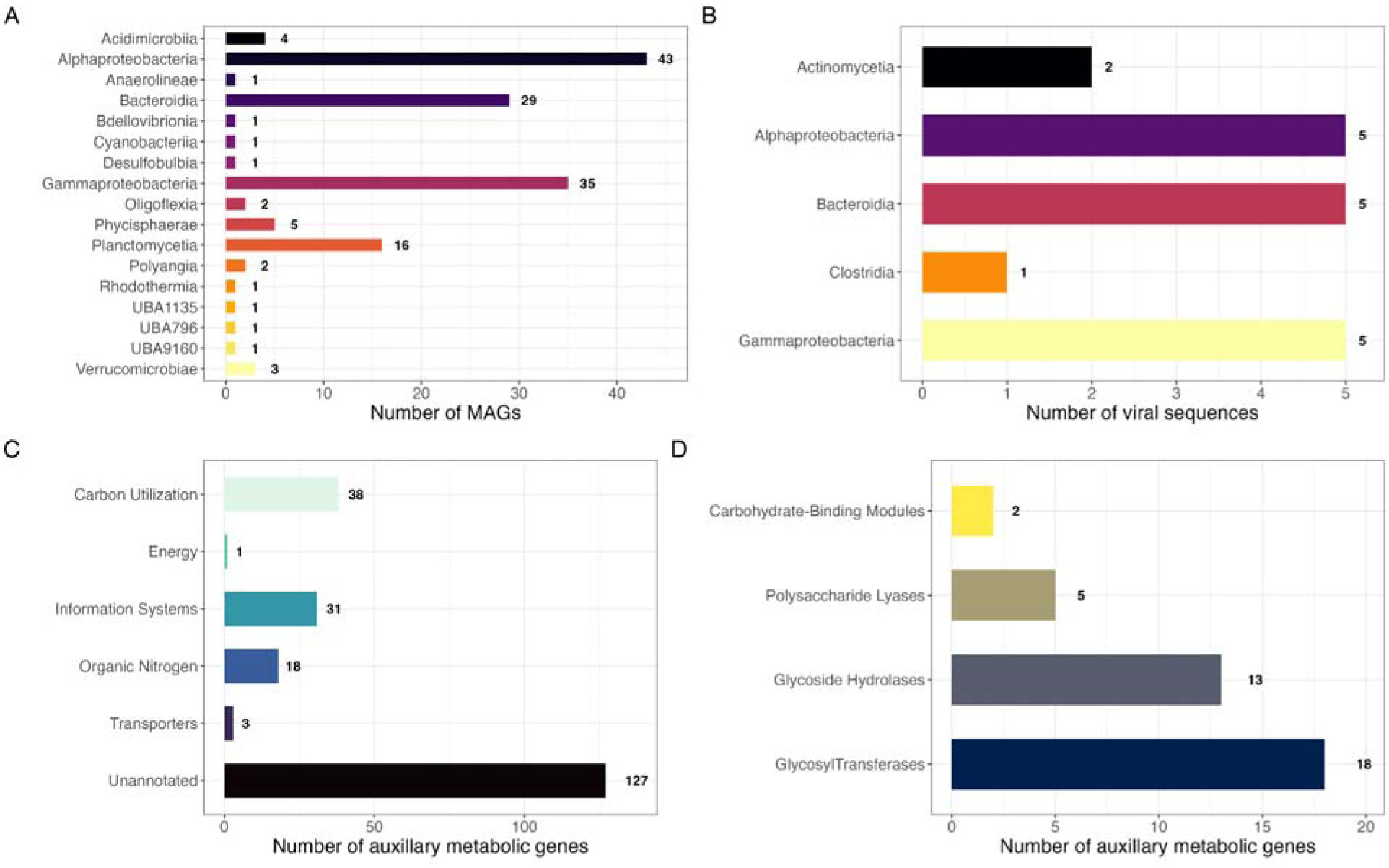
Viral-Bacterial interactions likely related to carbon metabolism. (A) Bar chart depicting the number of metagenome-assembled genomes (MAGs) identified in this study, colored by taxonomic class, and with the number to the right of the bar representing the total number of MAGs in each bar. (B) Bar chart depicting the viral host predictions, colored by host taxonomic class, and with the number to the right of the bar representing the total number of viral sequences in each bar. (C) Bar chart showing the number of predicted phage auxiliary metabolic genes (AMGs) summarized by DRAM-v distilled metabolic categories, with the number to the right of the bar representing the total number of AMGs in each bar. (D) Bar chart showing the number of predicted phage AMGs in different CAZyme families with-in the carbon utilization category, with the number to the right of the bar representing the total number of AMGs in each bar.

The taxonomic distribution of the recovered MAGs is broadly consistent with previous DNA-based surveys of *Zostera*-associated bacteria. These studies also identified Alphaproteobacteria, Gammaproteobacteria, and Bacteroidia as dominant classes, along with orders such as Rhodobacterales and Flavobacteriales [9–13]. Further a global study of *Z. marina* reported that Planctomycetia was enriched on leaves relative to surrounding water [8]. Together, these results suggest that this MAG collection complements previous DNA-based surveys, providing an opportunity for exploring deeper insight into the genomics and functional potential of abundant bacterial taxa within *Z. marina* ecosystems.

### Recovered phage largely predicted to infect most frequent MAG phyla

We used iPHoP [60] to predict bacteria-virus linkages but were only able to predict bacterial hosts for 18 vOTUs (Figure 2B). Of these, hosts generally were distributed across the most frequently recovered MAG classes (i.e., Alphaproteobacteria, Gammaproteobacteria, and Bacteroidia). While the MAG collection was added to the iPHoP database to enable prediction of direct links between MAGs and viruses, only one MAG, SGMAG-05 (a Saprospiraceae bacterium in the class Bacteroidia) was predicted to be host to a virus from the high-quality catalog. This phage, vOTU566, was identified as a prophage belonging to the Caudoviricetes. The remaining host-virus predictions were derived from iPHoP’s default genome library rather than the MAG collection. The limited number of host-virus links identified between the MAG and viral collections suggests that much more work remains to uncover these interactions within seagrass ecosystems. Future studies should endeavor to sequence the bacterial community alongside deeper virome sequencing to better capture viral diversity. Additionally, employing physical linking techniques such as Hi-C may help characterize novel host-viral interactions.

### Auxiliary metabolic genes point to phage role in carbon cycling

We used DRAM-v [55] to explore predicted AMG functions in the viral collection (Figure 2C). Over half of putative AMGs were unannotated (57.47%), similar to other recent studies of phage from the environment [71,72]. Given the importance of nitrogen and sulfur cycling in seagrass ecosystems, we searched the annotated AMGs for putative functions related to these processes. However no annotated AMGs had predictions related to nitrogen fixation or sulfur cycling. This absence could be biological (e.g., such genes may be more likely to be present in root or rhizosphere samples vs. leaves) or could reflect the limitations of recovering viruses from metagenomic datasets. Virome sequencing will likely be necessary to confirm whether viruses play a role in these nutrient cycles.

In contrast, we identified a variety of predicted AMGs with putative annotations of functions related to carbon utilization, particularly carbohydrate-active enzymes (CAZymes) involved in organic carbon cycling, sugar processing and plant degradation (Figure 2D). Seagrass beds are known as hot spots for carbon sequestration and key contributors to blue carbon storage [2,4]. Additionally previous work in the Mediterranean seagrass *Posidonia oceanica* has shown that rhizosphere sediments are enriched in sugars [46]. These findings may suggest a possible role for viruses in carbon cycling in seagrass beds, which may have important implications for blue carbon management and climate change mitigation.

## Conclusion

This study provides a valuable community resource and catalog of refined seagrass-associated bacterial and viral genomes, serving as a genomic foundation for future research. These comprehensive collections likely capture the most abundant groups associated with the leaves of *Z. marina*. While we found no evidence of AMGs related to nitrogen or sulfur metabolism, we instead report the presence of AMGs putatively linked to carbon utilization, particularly a diverse array of CAZymes that may be involved in organic carbon cycling. These results suggest that viruses may play an important role in carbon sequestration and blue carbon storage within seagrass ecosystems, which may have significant implications for climate change mitigation. Moving forward, research should prioritize investigating the role of phages through viromic techniques, which would offer deeper insights into viral ecology and phage contributions to carbon cycling in seagrass habitats.

## Declarations

### Ethics approval and consent to participate

Not applicable.

### Consent for publication

Not applicable.

### Availability of data and material

The raw metagenomic sequencing data was deposited at GenBank under accession no. <u>PRJNA1140276</u>. MAGs with > 90% completion estimates were deposited on NCBI under this BioProject at GenBank under accession no. <u>PRJNA1140276</u>, while MAGs with completion estimates between 80 and 90% were deposited on Zenodo (DOI: https://doi.org/10.5281/zenodo.14225973). Viral genomes were also deposited on Zenodo (DOI: https://doi.org/10.5281/zenodo.14226038). All code used in this work has been deposited on Github (<u>casett/Seagrass_MAG_and_Virus_Catalog</u>) and archived in Zenodo (DOI: https://doi.org/10.5281/zenodo.14226514).

### Authors’ Contributions

CLE conceived and designed the experiments, performed sampling, analyzed the data, prepared figures and/or tables, wrote and reviewed drafts of the paper. JES reviewed drafts of the paper.

## Acknowledgements

We would like to thank Jonathan A. Eisen for feedback on this manuscript. We would like to thank Katherine Dynarski (<u>ORCID: 0000-0001-5101-9666</u>) and Sonia Ghose (<u>ORCID: 0000-0001-5667-6876</u>) for their help with sample collection. We would also like to thank John J. Stachowicz for use of his scientific sampling permit, California Department of Fish and Wildlife Scientific Collecting Permit # SC 4874.

