## Supplemental Figures and Table Legends for "A genomic resource for exploring bacterial-viral dynamics in seagrass ecosystems"

bacteria, carbon cycling, metagenome-assembled genomes

**Supplemental Table Legends and Figures:**

**Table S1.** Metagenomic samples used for viral identification. Here we provide details on each metagenomic sample obtained from NCBI including NCBI accession number, seagrass species, sample type, collection location, data type, number of sequenced bases (in gigabases) reported in the Sequence Read Archive and appropriate data citation.

**Table S2.** Viral genome statistics. We identified 44 complete, 28 high-quality and 282 medium-quality draft viral genomes associated with seagrass ecosystems. Here, we list their proposed taxonomy, report on various genome and annotation statistics, and provide the NCBI Zenodo DOI for the dataset.

**Table S3.** Metagenome-assembled genome assembly statistics. We identified 85 high-quality and 62 medium-quality draft bacterial MAGs associated with *Z. marina* leaves. Here, we list their proposed taxonomy, report on various genome and annotation quality statistics, and provide the NCBI accession or Zenodo DOI for each MAG.

**Figure S1.** Viral catalog comparison to reference databases. (A) Gene-sharing network of vOTUs that formed viral clusters (VCs), including VCs unique to this dataset. Each node is a viral sequence and is colored by whether the sequence represents a viral sequence from this study or from reference data. Edges connect sequences with overlap in predicted protein content. (B) Bar chart displaying the number of viral sequences from this study assigned to viral classes based on geNomad. Bars are colored by viral class, and the number to the right of the bar represents the total number of sequences it represents.

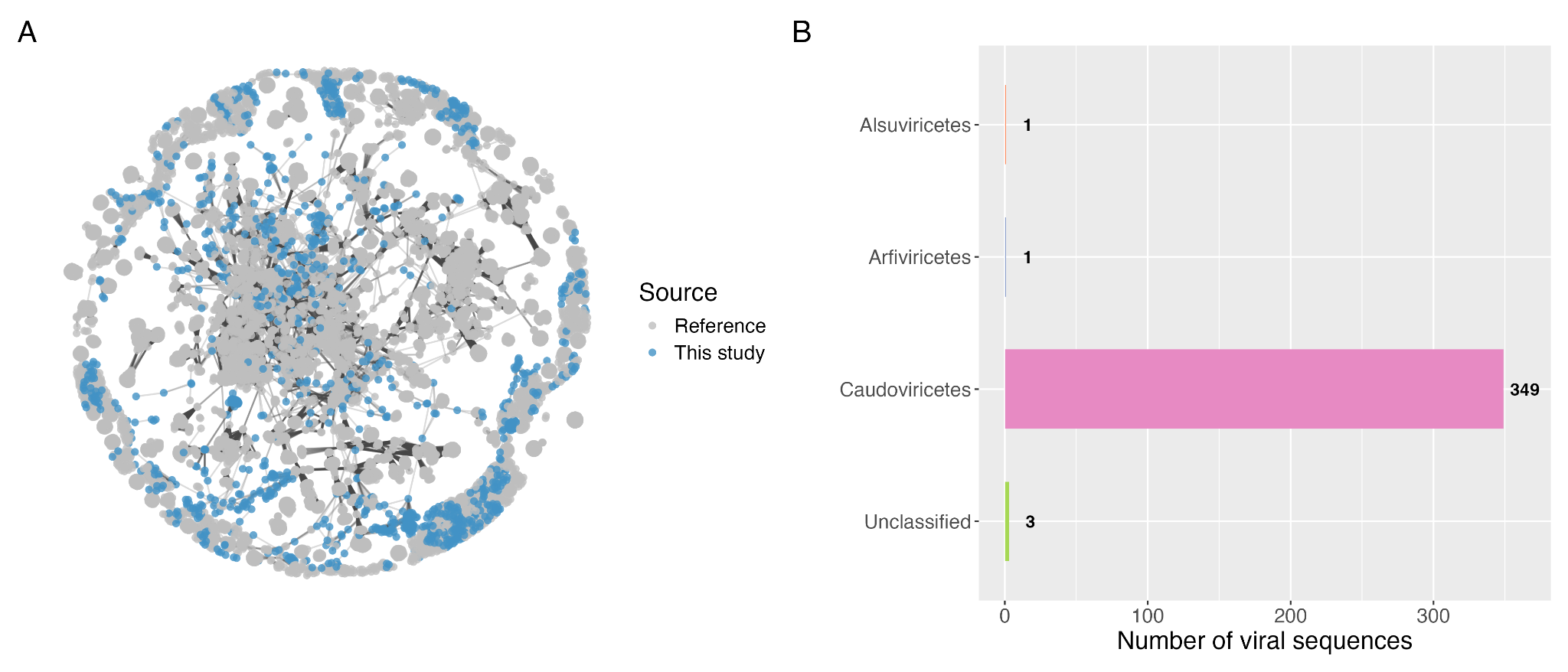
